# Mutagenesis of the Orco Odorant Receptor Co-receptor Impairs Olfactory Function in the Malaria Vector *Anopheles coluzzii*

**DOI:** 10.1101/2020.09.04.283424

**Authors:** Huahua Sun, Feng Liu, Zi Ye, Adam Baker, Laurence J. Zwiebel

## Abstract

Mosquitoes rely heavily on their olfactory systems for host seeking, selection of oviposition sites, and avoiding predators and other environmental dangers. Of these behaviors, the preferential selection of a human blood-meal host drives the vectorial capacity of anthropophilic female *Anopheles coluzzii* mosquitoes. Olfactory receptor neurons (ORNs) are dispersed across several appendages on the head and express an obligate odorant receptor co-receptor (Orco) coupled with a “tuning” odorant receptor (OR) to form heteromeric, odor-gated ion channels in the membrane of these neurons. To examine the mechanistic and functional contributions of Orco/OR complexes to the chemosensory processes of *An. coluzzii*, we utilized CRISPR/Cas9 gene editing to create a line of homozygous, Orco-knockout, mutant mosquitoes. As expected, *orco*^*-/-*^ ORNs across both adult and larval stages of *An. coluzzii* display significantly lower background activity and lack nearly all odor-evoked responses. In addition, blood-meal-seeking, adult female, *orco*^*- /-*^ mutant mosquitoes exhibit severely reduced attraction to human- and non-human-derived odors while gravid females are significantly less responsive to established oviposition attractants. These results reinforce observations in other insects that Orco is crucial in maintaining the activity of ORNs. In that light, it significantly influences a range of olfactory-driven behaviors central to the anthropophilic host preference that is critical to the vectorial capacity of *An. coluzzii* as a primary vector for human malaria.

## Introduction

As one of the most important vectors for the malaria parasite *Plasmodium falciparum*, which infected over 200 million people worldwide in 2018 and accounted for more than 400,000 deaths (WHO, 2019), the *Anopheles coluzzii* mosquito has long been a formidable concern to public health. As is the case for many hematophagous mosquito vectors that require human or animal blood for egg development and, hence, reproduction, blood feeding itself is responsible for transmitting *P. falciparium* and other pathogens between mosquito vectors and host animals (Aly et al., 2009; Beier, 1998; Whitten et al., 2006). The ability of *An. coluzzii* to vector human malaria is significantly enhanced by its distinctive anthropophilic host preference (Montell & Zwiebel, 2016), which involves a broad array of sensory cues, including heat, CO_2_, and odors released from human emanations, to locate and select blood-meal hosts (Takken & Knols, 1999). Indeed, human-derived odors have been shown to activate a spectrum of Anopheline chemosensory neurons across antennae, maxillary palps and labella (Lu et al., 2007; Qiu et al., 2006; Kwon et al., 2006).

Several types of chemosensory receptors, such as odorant receptors (ORs), ionotropic receptors (IRs), and gustatory receptors (GRs), are expressed in the dendrite membranes of diverse classes of chemosensory neurons to drive the generation of stimulus-evoked action potentials (Suh et al., 2014). Olfactory receptor neurons (ORNs), which are housed in diverse classes of hair-like sensilla dispersed across the antennae, maxillary palps and proboscis, express ORs that are responsible for detecting a range of general and specialized odors (reviewed in (Suh et al., 2014)). Insect ORs comprise very large gene families that are generally characterized by highly divergent “tuning” receptors with relatively low sequence similarity across different taxa of insects that are responsible for the recognition and binding of odorants (Hansson & Stensmyr, 2011). The notable exception to that OR conservation paradigm is the odorant receptor co-receptor (Orco) gene that is highly conserved and has been shown to be orthologous among various insect species (W. D. Jones et al., 2005). More importantly, Orco plays an obligate role in establishing functional insect ORNs, as it has been shown to couple with discrete tuning ORs to form heteromeric complexes required for dendritic membrane trafficking, odorant recognition and olfactory signal transduction (R. Benton, 2006; Larsson et al., 2004). Unlike mammalian ORs, which function as GPCRs, insect OR complexes act as ligand-gated ion channels *in vivo* (Wicher et al., 2009; Sato et al., 2008) as well as *in vitro* in numerous heterologous expression platforms (Hallem and Carlson, 2006; Wang *et al*., 2010). Pharmacological (Jones *et al*., 2011; Pask PLosONe 2011+Kumar etal 2013) and, more recently, structural studies of Orco complexes have further illuminated the centrality of the Orco/Or ion channel complex in insect olfactory signal transduction (Butterwick et al., 2018).

In light of its central role in insect olfactory transduction, several studies have used various gene-silencing/targeting strategies to generate *orco* mutants. In studies across insect species, such as the vinegar fly (*Drosophila melanogaster*), Dengue mosquito (*Aedes aegypti*), hawkmoth (*Manduca sexta*), clonal raider ant (*Ooceraea biroi*), ponerine ant (*Harpegnathos saltator*) and the migratory locust (*Locusta migratoria*), OR-based olfactory signaling is either completely silenced or dramatically impaired when Orco functionality is knocked out (Larsson et al., 2004; Degennaro et al., 2013; Li et al., 2016; Guo et al., 2017; Yan et al., 2017; Fandino et al., 2019; Trible et al., 2017). To further our understanding of the olfactory physiology as well as promote novel malaria control strategies for *An. coluzzii*, we have used CRISPR/Cas9 gene editing to generate the first Orco knockout line in Anopheline mosquitoes. An examination of olfactory physiology in both adults and larvae reveals that *orco*^-/-^ *An. coluzzii* mutants are similarly unable to detect a majority of general odorants in the OR-expressing ORNs throughout their life cycle. Taken together, these data reinforce the requirement of Orco functionality in Anopheline olfaction and vectorial capacity, thereby highlighting Orco as a critical molecular target for malaria control.

## Materials and Methods

### Mosquito maintenance

*An. coluzzii* (SUA 2La/2La), previously known as *Anopheles gambiae sensu stricto* “M-form”(Coetzee et al., 2013), originated from Suakoko, Liberia, were reared using previously described protocols (Fox et al., 2001; Y. T. Qiu et al., 2004). All mosquito lines were reared at 27°C, 75% relative humidity under a 12:12 light:dark cycle and supplied with 10% sugar water in the Vanderbilt University Insectary (Fox et al., 2001; Suh et al., 2016). Mosquito larvae were reared in distilled water with approximately 300 larvae per rearing pan in 1L H_2_O. Larval food was prepared by dissolving 0.12g/mL Kaytee Koi’s Choice premium fish food (Chilton, WI, US) and 0.06g/mL yeast in distilled water and incubating at 4 °C overnight for fermentation. For 1^st^ and 2^nd^ instar larvae, 0.08mL larval food solution was added daily into each rearing pan. For 3^rd^ and 4^th^ instar larvae, 1mL larval food solution was added to each pan daily.

### CRISPR-Cas9 gene editing

The CRISPR-Cas9 gene editing procedure was carried out as previously described (Liu et al., 2020), with minor modifications. The CRISPR gene targeting vector was a generous gift from the lab of Dr. Andrea Crisanti of Imperial College London, UK (Hammond et al., 2016). The single guide RNA (sgRNA) sequences for Orco (AGAP002560) were designed for high efficiency using the CHOPCHOP online tool (http://chopchop.cbu.uib.no/), commercially synthesized (Integrated DNA Technologies, Coralville, IA) and subcloned into the CRISPR vector via Golden Gate cloning (New England Biolabs, Ipswich, MA) (Table S1). The homology templates were constructed based on a pHD-DsRed vector (a gift from Kate O’Connor-Giles; Addgene plasmid #51434; http://n2t.net/addgene:51434; RRID: Addgene 51434). The 2-kb homology arms extending in either direction from the double-stranded break (DSB) sites were PCR amplified and sequentially inserted into the AarI/ SapI restriction sites on the vector (Table S1).

The microinjection protocol was carried out as described (Pondeville et al., 2014; Ye et al., 2020). Briefly, newly laid (approximately 1h-old) embryos of wild-type *An. coluzzii* were immediately collected and aligned on a filter paper moistened with 25mM sodium chloride solution. All the embryos were fixed on a coverslip using double-sided tape, and a drop of halocarbon oil 27 was applied to cover the embryos. The coverslip was further fixed on a slide under a Zeiss Axiovert 35 microscope with a 40× objective. The microinjection was performed using an Eppendorf FemtoJet 5247 (Eppendorf, Enfield, CT) and quartz needles prepared using a customized protocol (Sutter Instrument, Novato, CA). The gene targeting vector and the homology template were diluted to 300ng/μL each and co-injected to maximum capacity for each embryo. Injected embryos were subsequently placed in deionized water with artificial sea salt (0.3g/L) and thereafter reared under normal VU insectary conditions.

The first generation (G0) of injected adults were separated based on gender and crossed to 5× wild-type gender counterparts. Their offspring (F1) were manually screened for DsRed-derived red eye fluorescence using an Olympus BX60 Compound Fluorescent Microscope (Olympus, PA). Red-eyed F1 males were individually crossed to wild-type females for 5 generations to establish a heterozygotic line. Genomic DNA extractions using Qiagen Gel Extraction protocols (Qiagen, Germantown, MD) and PCR analyses were carried after the individuals were allowed to mate to validate the fluorescence marker insertion using primers that cover DSB sites (Table S1). PCR products were sequenced to confirm the accuracy of the genomic insertion. Validated heterozygous mutant lines were thereafter back-crossed to wild-type *An. coluzzii* for 5 generations before putative homozygous individuals were manually screened for DsRed-derived red-eye fluorescence intensity. Putative homozygous mutant individuals were mated to each other before being sacrificed for genomic DNA extraction and PCR analyses (as above) to confirm their genotypes.

### Immunohistochemistry

Immunofluorescence was performed as described, with minor modifications (Kwon et al., 2006; Pitts et al., 2004). The antennae and maxillary palps were hand dissected and prefixed in a solution of 4% paraformaldehyde in PBST (137mM NaCl, 2.7mM KCl, 10mM Na_2_HPO_4_, 2mM KH_2_PO_4_ and 0.1% Triton X-100, pH 7.5) on ice for 30min. Samples were rinsed three times with PBST for 5 min on ice and then embedded and frozen in Tissue Freezing Medium (Electron Microscopy Sciences, PA). Samples were sectioned (8μm) using a CM1900 cryostat (Leica Microsystems, Bannockburn, IL) at -20°C, and sections were collected on Superfrost plus slides (VWR Scientific, Radnor, PA). Slides were dried at room temperature (RT) for 30 min, fixed in PBST solution containing 4% paraformaldehyde for 15 min, and then rinsed 3× in PBST for 5 min. Slides were blocked using 5% Normal Goat Serum (NGS, Sigma-Aldrich, St. Louis, MO) in PBST in the dark at RT for 1h, in covered HybriWell sealing chambers (Grace Bio-Labs, Bend, OR).

Primary Rabbit α-Orco antisera (Pitts et al., 2004) were diluted 1:2000 in 5% NGS/PBST and applied onto the slides and incubated at 4°C overnight. Sequentially, slides were rinsed 3× in PBST for 5min and incubated with secondary antibody, Goat α-Rabbit-Cy3 (Jackson ImmunoResearch, West Grove, PA) at a 1:2000 dilution in 5% NGS/PBST at RT for 2h and then rinsed, as above. Nuclei were labeled with 300nM DAPI (Invitrogen, Carlsbad, CA, USA). After washing 3× in PBST, the slides were mounted in Vectashield fluorescent medium (Vector Laboratories, Burlingame, CA). An Olympus FV-1000 confocal microscope equipped with a 100× oil immersion objective (Vanderbilt University Cell Imaging Shared Resource Core) was used to collect images (1024*1024-pixel resolution). Laser wavelengths of 405nm and 543nm were used to detect DAPI and Cy3, respectively.

### Electrophysiology

Electroantennograms (EAGs) were conducted using non-blood-fed, 5- to 7-day-old, mated adult females according to (Suh et al., 2016), with modifications. Here, the mosquito head was dissected, fixed to a glass microscope slide, and the terminal flagellar segment of each antennae was transected. The decapitated head with transected antennae was subsequently connected to a glass recording electrode filled with Ringer solution (96mM NaCl, 2mM KCl, 1mM MgCl_2_, 1mM CaCl_2_, 5mM HEPES, pH = 7.5), in which a AgCl-coated sliver wire was placed in contact to complete a circuit with a reference electrode inserted into the back of the head. Antennal preparations were continuously exposed to a humidified, charcoal-filtered air flow (1.84L/min) transferred through a borosilicate glass tube (inner diameter = 0.8cm) that was exposed to the preparation at a distance of 5mm. Stimulus cartridges were prepared by transferring 10μl of test or control stimuli solutions to filter paper (3×50mm), which was then placed inside a 6-inch Pasteur pipette. Odorant stimuli were delivered to antennal preparations for 500ms through a hole placed on the side of the glass tube located 10cm from the open end of the delivery tube (1.08L/min), where it was mixed with the continuous air flow using a dedicated stimulus controller (Syntech, Hilversum, The Netherlands). A charcoal-filtered air flow (0.76L/min) was simultaneously delivered from another valve through a blank pipette into the glass tube at the same distance from the preparation in order to minimize changes in flow rate during odor stimulation. The resulting signals were amplified 10× and imported into a PC via an intelligent data acquisition controller (IDAC, Syntech, Hilversum, The Netherlands) interface box, and the recordings were analyzed offline using EAG software (EAG Version 2.7, Syntech, Hilversum, The Netherlands). Maximal response amplitudes of each test stimuli were normalized after dividing by the control (solvent alone) responses.

Adult single sensillum recordings (SSRs) were carried out as previously described (Liu et al., 2020), with minor modifications. Electrophysiological recordings on 3^rd^ instar larval antennal sensory cones were carried out according to (Sun et al., 2020). Specifically, mosquitoes (adult or larva) were mounted on glass microscope slides (76×26mm) (Ghaninia et al., 2007) and their antennae were fixed using double-sided tape to a cover slip resting on a small bead of dental wax to facilitate manipulation. The cover slip was placed at approximately 30 degrees to the mosquito head. Once mounted, the specimen was placed in an Olympus BX51WI microscope and antennae were viewed at high magnification (1000×). Two tungsten microelectrodes were sharpened in 10% KNO2 at 10V. The grounded reference electrode was inserted into the compound eye of the mosquito/larvae using a WPI micromanipulator, and the recording electrode was connected to the pre-amplifier (Syntech universal AC/DC 10x, Syntech, Hilversum, The Netherlands) and inserted into the shaft of the olfactory sensillum to complete the electrical circuit to record ORN potentials extracellularly (Den Otter et al., 1980). Controlled manipulation of the recording electrode was performed using a Burleigh micromanipulator (Model PCS6000). The preamplifier was connected to an analog-to-digital signal converter (IDAC-4, Syntech, Hilversum, The Netherlands), which in turn was connected to a PC for signal recording and visualization.

Odorant compounds of the highest available purity, typically ≧99% (Sigma-Aldrich), were diluted in paraffin oil to make 1% v/v (liquids) or m/v (solids) solutions. For each compound, a 10-μL portion was dispersed onto filter paper (3×10mm), which was then inserted into a 6-inch Pasteur pipette to create a stimulus cartridge. A sample containing the solvent alone served as the control. The airflow across the antennae was maintained at a constant 20mL/s throughout the experiment. Purified and humidified air was delivered to the preparation through a glass tube (10-mm inner diameter) perforated by a small hole 10cm away from the end of the tube into which the tip of the Pasteur pipette stimulus cartridge could be inserted. The stimulus was delivered to the sensilla by inserting the tip of the stimulus cartridge into this hole and diverting a portion of the air stream (0.5L/min) through the stimulus cartridge for 500ms using a Syntech stimulus controller CS-55 (Syntech, Hilversum, The Netherlands). The distance between the end of the glass tube and the antennae was ∼1cm. Signals were recorded for 10s, starting 1sbefore stimulation, and the action potentials were transformed off-line into spikes using AutoSpike (Syntech, Hilversum, The Netherlands). Spikes were counted across a single 0.5-s pre-stimulus interval as well as across four 0.5-s post-stimulus windows. “Pre-stimulus” is defined as the time immediately preceding odorant delivery and “Post-stimulus” as the time immediately following the onset of neuronal activation. Normalized neuronal responses were calculated by subtracting the pre-stimulus activity from the spike number observed during 0.5-s post-stimulus measurement and thereafter subtracting similarly normalized solvent responses. Firing frequency was counted in units of spikes per second.

### Human host proximity assay

Human host proximity bioassays were similar to that described in DeGennaro et al. (2013). For each trial, 50-60 5-to 7-day-old, wild-type and mutant adult female mosquitoes were transferred into a Bugdorm® insect rearing cage (polypropylene, 30×30×30cm) and kept under fasting conditions (i.e. with access only to water) for 16-24h prior to each trial. Two human volunteers (who had used no scented products for 48h and had not washed their hands for 4h prior to experiment) each wore an odor-free laboratory glove for 30min to induce perspiration; the glove was then removed and the volunteer’s hand was immediately inserted into a 24-oz. Solo cup (Dart Container Corp, MI) with the bottom removed. The bottom opening of the cup was held tightly with odor-free dental wax (Orthomechanic, Broken Arrow, OK) to the mesh side of the cage. Within the cup, each hand was held 2.5cm from the mesh side of the cage, to prevent mosquitoes from making direct contact. A GoPro camera (GoPro, CA) was positioned to take images of mosquitoes responding to the hand (or to the empty control). Trials ran for 5min and images were acquired digitally for the entire period. To quantify mosquito responses, the number of mosquitoes landing on the cup bottom were manually counted during the entire assay window. Normalized mosquito landing scores were generated by taking the total number of landings and dividing by the total number of mosquitoes in the cage.

### Non-human Dual Choice Preference Assay

Mutant and wild-type mosquitoes were examined in a dual-choice assay, with minor modifications (Wondwosen et al., 2016), using Limburger cheese (St. Mang, Bavaria) as a general Anopheline attractant (Knols & De Jong, 1996; Owino, 2011). A two-port, still-air olfactometer, constructed by modifying a Bugdorm® insect rearing cage, was used to test mosquito attraction preference for the cheese volatiles. For each replicate, 50-60 5-to 7-day-old, mated adult females were allowed to acclimatize for 10min in a 64-oz Solo cup (Dart Container Corp, MI) covered with clear vinyl for easy viewing. Following this, two air-activated heat sources (HotHands^®^, Heat Max, Dalton, GA) (Saveer et al., 2018) were simultaneously introduced into opposing, cylindrical, vinyl mesh-covered arms (13 cm × 9cm; L: d), positioned at opposite ends of the Bugdorm® insect rearing cage, together with 5g Limburger cheese which was placed on top of the heat source. Immediately after placement of the cheese, the mesh on top of the mosquito cup was removed, releasing mosquitoes and allowing them to enter either of the cylinders (with or without cheese). This assay ran for 15min, after which the number of mosquitoes trapped in each cylinder was counted. A preference index was calculated by dividing the total number of mosquitoes in each cylinder by the total mosquitoes released.

### Oviposition preference assay

The oviposition preference assay was carried out for both wild-type and mutant mosquitoes in a dual-choice assay in a Bugdorm® insect rearing cage, as previously described (Saveer et al., 2018), with minor modifications. The oviposition blends were comprised of seven compounds (2-n-propylphenol, 2-methylphenol, 4-methylphenol, 4-ethylphenol, nonanal, indole, and skatole) each at 10^−8^ M in ethanol (Choo et al., 2015, 2018; Himeidan et al., 2013; Hughes et al., 2010; Rinker et al., 2013; Zhu et al., 2013). Briefly, two plastic egg cups (top opening = 7.0cm, height = 6.0cm, bottom = 5.0cm) filled with 10mL dH2O as an oviposition substrate were placed at opposite corners of the assay cage. A pair of borosilicate glass vials (14.65×19 mm; Qorpak, Bridgeville, PA) with screen tops (10×10mm) filled with 1mL oviposition blend or solvent control were placed inside the egg cup (Fig. 5B). Experiments were initiated approximately 1 h prior to scotophase (Zeitgeber time 11) by introducing 20 gravid females (48-h post-blood fed) into the assay cage, and the total number of eggs in the two oviposition cups were manually counted on the following day (ZT9). Five bioassay cages were prepared for every treatment and control (solvent-solvent) trial.

## Results

### Generation of *An. coluzzii orco* mutant lines

In order to generate a loss-of-function *An. coluzzii* mutant line for the *orco* gene, guide RNAs (gRNA) were designed using the CHOPCHOP online tool (Labun et al., 2019) to target the third exon to generate a truncated, nonfunctional protein (Figure 1A, B). At the same time, a DsRed visible eye color marker was inserted into the DSB site of the *orco* gene target via homologous recombination to drive the expression of red eye color in mutant mosquitoes to facilitate selection (Figure1C). Heterozygous (*orco*^*+/-*^) mutant mosquito lines were thereafter back crossed to wild-type *An. coluzzii* for five generations before homozygous (*orco*^*-/-*^) mutant lines were established by self-crossing. Using PCR with primers designed to amplify genomic DNA sequences covering the DSB site, each mutant line was molecularly confirmed as having homozygotic insertions into the *orco* gene; this also served to confirm the specific insertion of eye color marker (Figure 1D).

**Figure 1.**
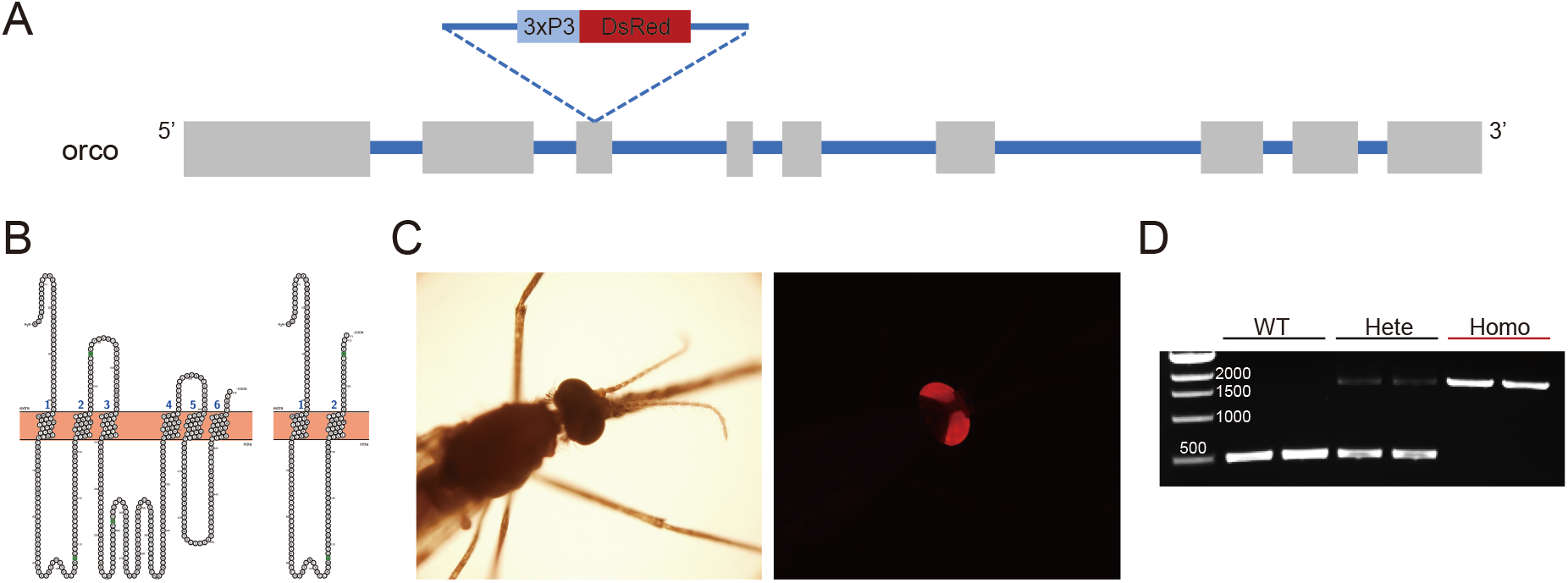
Generation of orco mutants using CRISPR-Cas9 gene editing. (A) Schematic of the organization of the *An. coluzzii orco* gene with exons depicted as boxes and introns as lines showing the gRNA target site and DsRed marker gene inserted into the third exon of *orco* gene. (B) Truncation of amino acid sequences after insertion of marker gene in the open reading frame of *orco* gene. Left: predicted model using online program (http://wlab.ethz.ch/protter) for secondary structure and membrane orientation of wild-type Orco protein; Right: predicted model for secondary structure of hypothetically truncated Orco protein in *orco*^*-/-*^ mutant. (C) DsRed florescence marker present in the compound eye of mutant mosquitoes with *orco* gene 3xP3 DsRed knockin insertion (left panel: phase contrast microscope, bright field under regular light; right panel: dark field under UV light). (D) Genotyping of the mutant mosquito lines using a pair of PCR primers that amplify across the DSB site in exon 3 of the *An. coluzzii* orco gene.

To confirm the absence of the Orco protein in *orco*^-/-^ ORNs, the antennae and maxillary palps from wild-type and mutant female *An. coluzzii* were hand dissected and immunohistochemically stained using α-Orco antibodies. Examination of appendages from wild-type *An. coluzzii* females confirmed that Orco is robustly expressed in both the antennae (Figure 2A, B) and the maxillary palps (Figure E and F); in the *orco*^-/-^, Orco-positive cells were not detectable in either antennae (Figure 2C, D) nor maxillary palps (Figure G and H). Taken together, these results strongly support our hypothesis that *Orco* expression is indeed disrupted and the protein is absent in *orco*^-/-^ ORNs.

**Figure 2.**
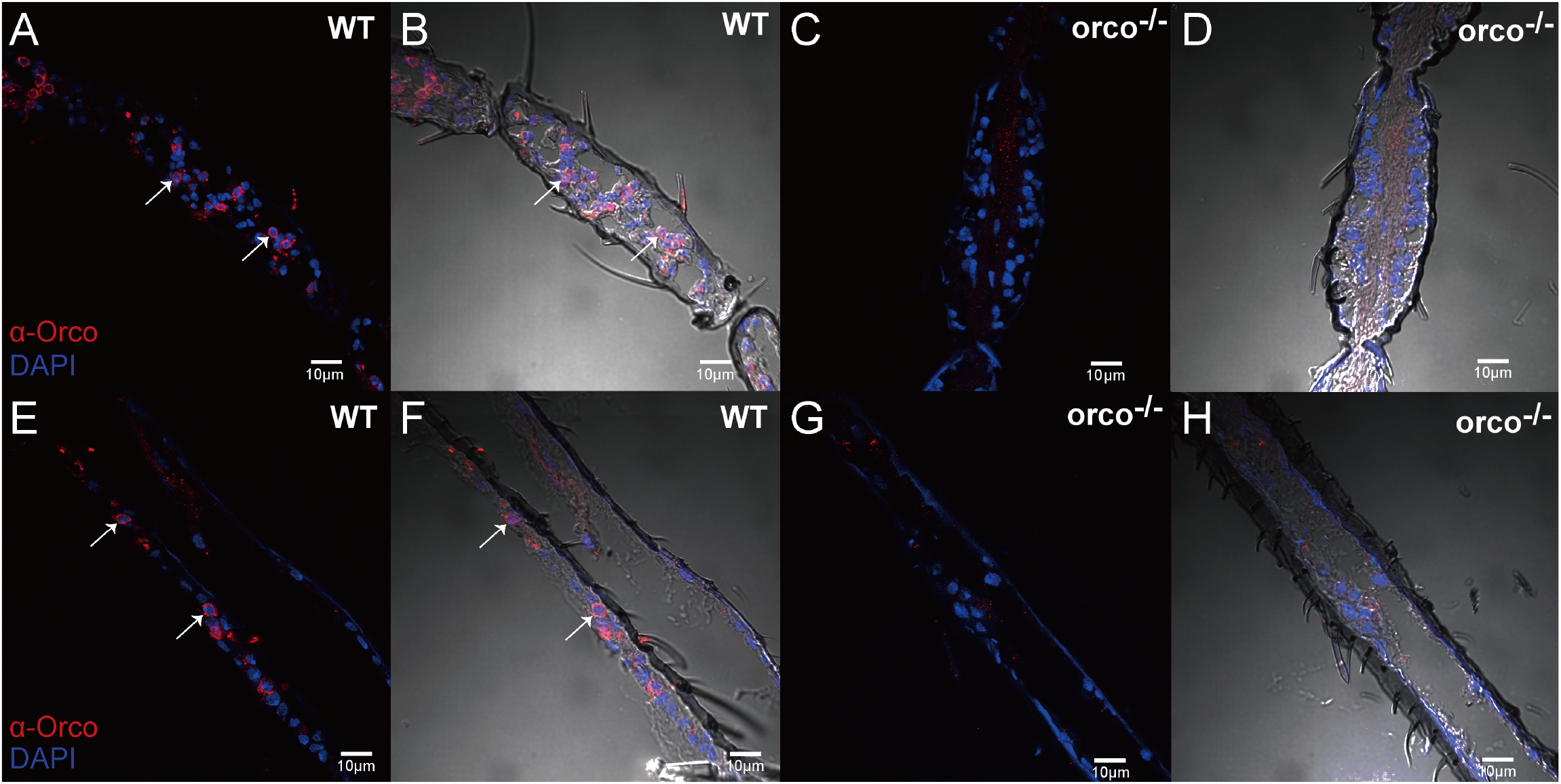
Localization of Orco on *An. coluzzii* antennae and maxillary palp. (A, B) Immunostaining with α-Orco antibodies indicating Orco-positive cells in the female wild-type mosquito antenna (red); (C, D) Immunostaining with α-Orco antibodies of the antennae from female *orco*^*-/-*^ mutant is devoid of Orco protein; (E, F) Immunostaining of Orco-positive cells on *An. coluzzii* female wild-type maxillary palp (red); (G, H) Immunostaining with α-Orco antibodies on *An. coluzzii* female *orco*^*-/-*^ mutant maxillary palp is devoid of Orco protein. Orco-positive cells are indicated by white arrows. Scale bars = 10μm.

### Reduced antennal responses to a broad panel of odors

To investigate the impact of the *orco*^-/-^ mutation on the peripheral responses across the *An. coluzzii* antennae, comparative EAGs were carried out on wild-type and mutant mosquitoes, using a broad odorant panel of 53 compounds ranging across 9 different chemical classes (Supplemental Table S2). As expected, wild-type mosquitoes displayed robust EAG responses to all of these chemical groups, while *orco*^-/-^ antennae were largely unresponsive when challenged with esters, ketones, alcohols, aldehydes, terpenes, sulfurs, and thiazoles (Figure 3A/B). This confirms the essential role that Orco plays in *An. coluzzii* and by extension other Anopheline malaria vectors in the detection of a wide range of odorants that comprise both host and environmental cues. Interestingly, no significant difference was observed between *orco*^-/-^ and wild-type EAGs in response to two amines (butylamine and 3-methylpiperidine) and two carboxylic acids (2-oxobutyric acid and L-(+)-lactic acid) from our panel (Figure 3B). The integrity of these responses in mutant antennae indicates that other classes of non-Orco/Or-dependent chemosensory receptors, most likely *An. coluzzii* IRs, must be involved in the detection of compounds in these chemical groups.

**Figure 3.**
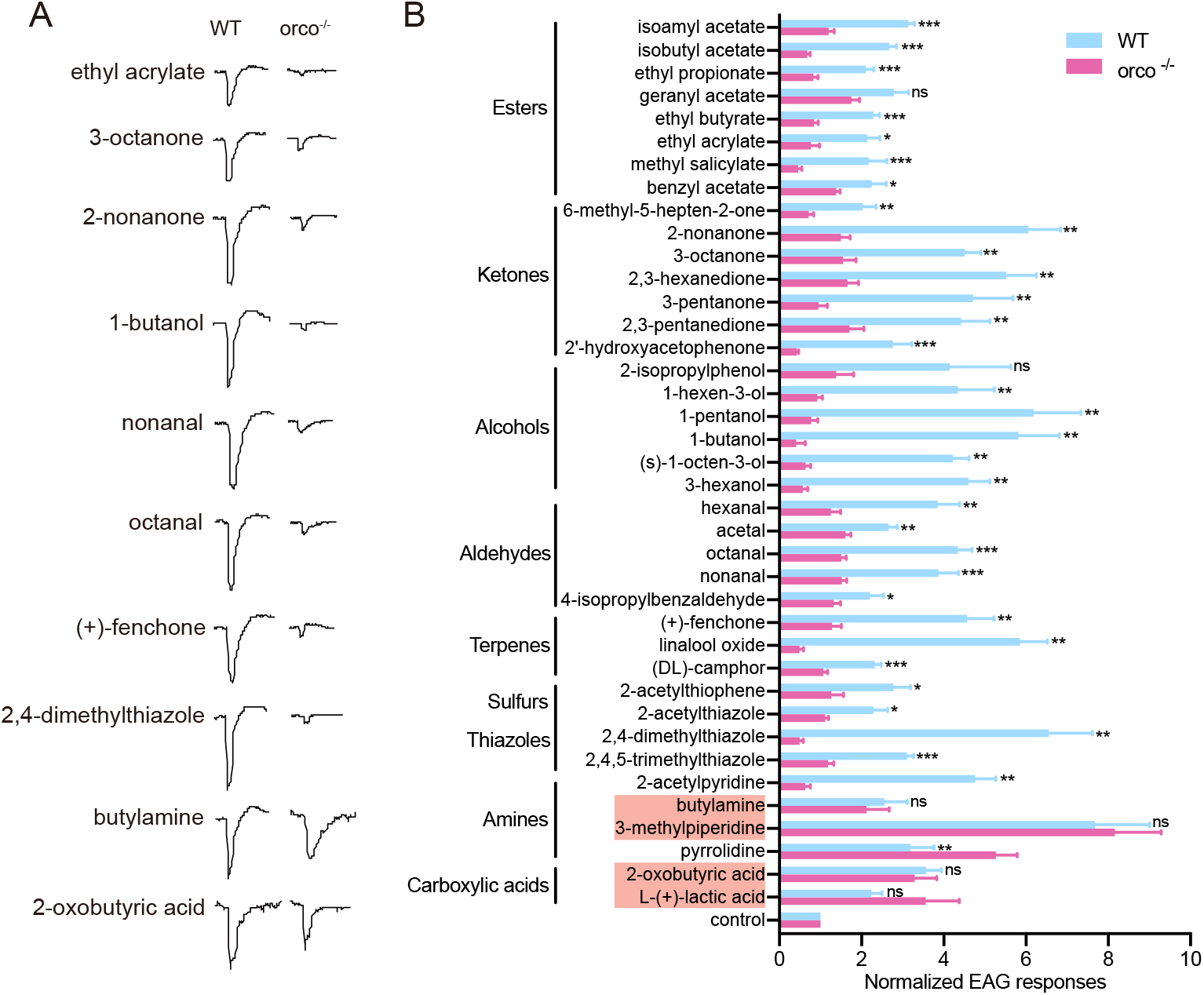
EAG responses of wild-type and *orco*^*-/-*^ *An. coluzzii* to a broad panel of odorants: (A) Representative EAG response trace of wild-type and *orco*^*-/-*^ mosquito to different odorants; (B) Comparison of EAG responses of wild-type and *orco*^*-/-*^ *An. coluzzii* to 53 odorants in different chemical classes (n=6). EAG responses were normalized to the solvent control (paraffin oil) for each odorant at a concentration of 10^−1^ dilution. Nonparametric Mann-Whitney test was applied in the statistical analysis, with p ≥0.05 indicating no significance (ns), and p<0.05 (*), p<0.01 (**) and p<0.001 (***) as significant differences. Two amines and two acids eliciting no significant difference between wild-type and *orco*^*-/-*^ mosquito are highlighted with red shadow.

### Selective insensitivity to odorants

As Orco is required to form a functional ion channel in the dendritic membrane (Sato *et al*., 2008; Butterwick *et al*., 2018), we hypothesized that OR-expressing ORNs in *An. coluzzii orco*^-/-^ mutants would show deficits in background activity and sensitivity to chemical agonists. Previous single sensillum electrophysiological recording (SSR) studies on maxillary palp capitate peg (cp) sensillum demonstrated that, in addition to the characteristic CO_2_ sensitivity of the large amplitude “A” neuron expressing a triad of GRs (F. Liu et al., 2020; Lu et al., 2007), there is a set of smaller amplitude responses derived from the “B/C” ORNs that function together with Orco in complex with either Or8 or Or28. The first of these, Or8, is extremely sensitive to several odorants associated with human sweat, most notably 1-octen-3-ol while Or28 has a broad odorant sensitivity (Lu *et al*., 2007). When compared with wild-type maxillary palp cp responses, in which the cpA as well as the cpB/C neurons retain normal background activity as well as robust responses to CO_2_ and 1-octen-3-ol, respectively (Figure 4A), the cpB/C neurons in *orco*^-/-^ mutants were almost completely silent and non-responsive to 1-octen-3-ol stimulation, with only rare residual spikes (Figure 4E and D). As expected, the cpA neuron showed no difference insofar as background activity or responses to CO_2_ when comparing wild-type and o*rco*^-/-^ cps (Figures 4C and B).

**Figure 4.**
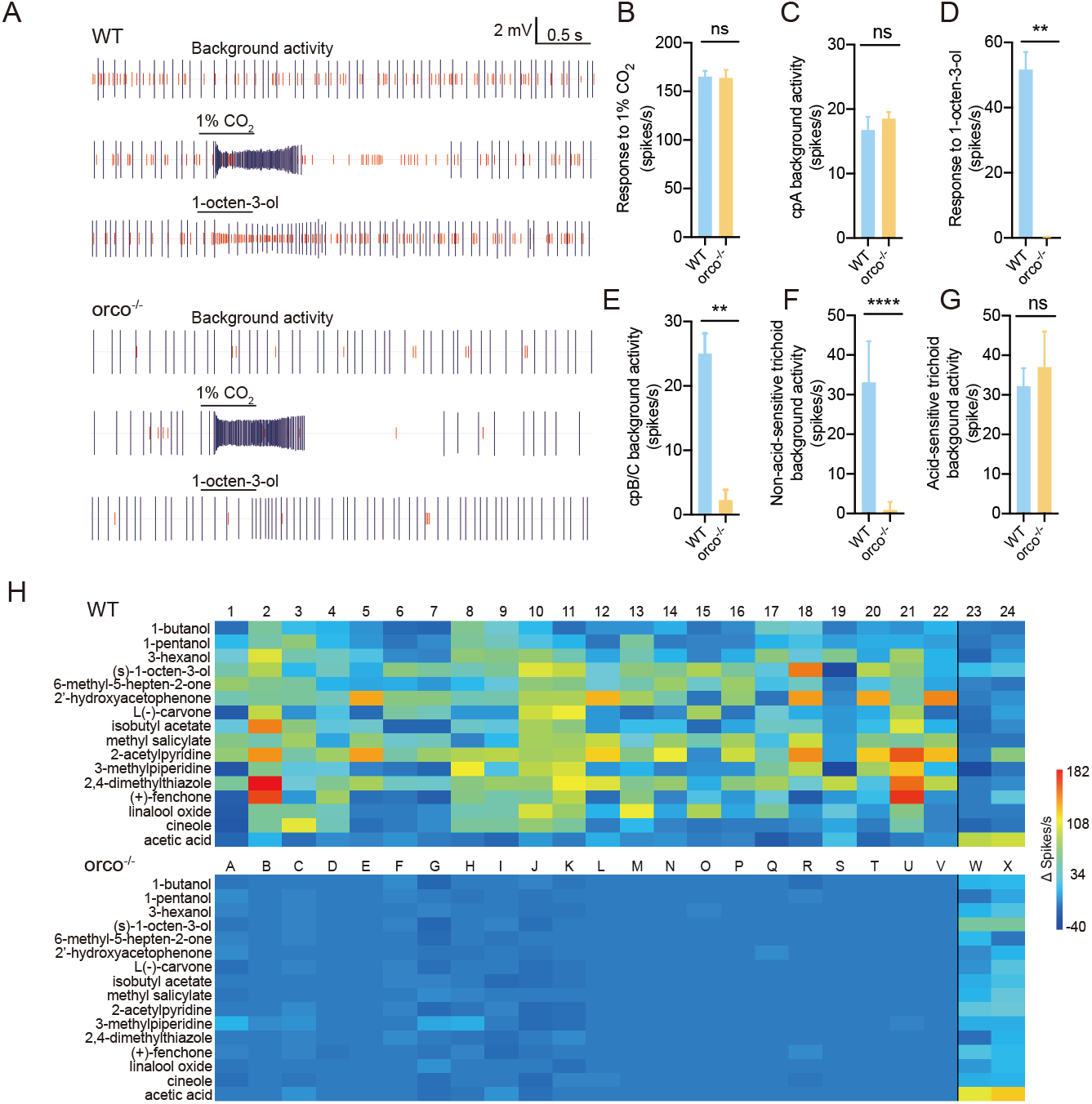
Responses of ORNs in maxillary palp and antenna of wild-type and orco^-/-^ mosquito to odorants: (A) Representative response traces of maxillary palp capitate peg sensillum to 1% CO_2_ and 1% 1-octen-3-ol in paraffin oil. Top: wild-type (WT), bottom: orco^-/-^. (B) Responses of maxillary palp cpA neurons to CO_2_ between wild-type and orco^-/-^ *An. coluzzii* (n=8); (C) Background activity of maxillary palp cpA neuron of wild-type and orco^-/-^ *An. coluzzii* (n=8); (D) Responses of maxillary palp cpB/C neuron to 1% 1-octen-3-ol in wild-type and orco^-/-^ *An. coluzzii* (n=8); (E) Background activity of maxillary palp cpB/C neurons of wild-type and orco^-/-^ *An. coluzzii* (n=8); (F) Background neuronal activity of a random survey across non-acid-sensitive antennal trichoid sensillum in wild-type and orco^-/-^ *An. coluzzii* (n=8); (G) Background neuronal activity of a random survey across acid-sensitive antennal trichoid sensillum in wild-type and orco^-/-^ *An. coluzzii* (n=2-5); (H) SSR responses to a 16 odorant subpanel of 24 individual trichoid sensilla randomly sampled across the antenna of wild-type (Top) and orco^-/-^ (Bottom) *An. coluzzii*. The black line segregates the first 22 individual trichoid sensilla, which display variable responses to one or more of the odorant cues but no response to acetic acid, and the last 2 individual trichoid sensilla, which were largely unresponsive to most of the stimulus panel while displaying strong responses to acetic acid. Nonparametric Mann-Whitney test was applied in all statistical analyses, with p<0.05 (*), p<0.01 (**) and p<0.001 (***) indicating significant differences.

While the maxillary palp, owing in large part to its uniform sensillar composition, is the best characterized sensory appendage of *An. coluzzii*, the majority of the OR-expressing ORNs reside in the highly heterogeneous population of trichoid sensillum on the mosquito antenna (Schultze et al., 2014; Pitts et al., 2004). In order to examine the effect of Orco deficiencies on that population of cells, we carried out a random sampling of antennal trichoid sensilla using SSR. Of the 24 wild-type trichoid sensilla sampled across multiple antennal flagella, all displayed significant background spiking (Figure 4F and G) while only 22 showed robust excitatory responses to at least one compound in a panel of 12 odorants (Figure 4H). The remaining two trichoid sensilla displayed relatively modest responses to the bulk of the odor panel along with selectively strong responses to acetic acid. When the same random sampling was carried out across *orco*^-/-^ mutant antennae, no odor-evoked responses were present in most trichoid sensilla, with the notable exception of acetic acid-sensitive trichoid sensilla (Figure 4H).

Interestingly, while some trichoid sensilla in the mutant mosquito present absolutely no background activity (Figure 4F), there are also some that show residual spikes, which nevertheless are insensitive to the same panel of odor stimulation used for wild-type mosquito (Supplementary Figure S1). Unsurprisingly, the acetic acid-sensitive sensilla still show the same strong response to acetic acid in *orco*^-/-^ preparations, suggesting Orco is not involved in detecting acetic acid (Supplementary Figure S2). As previously reported in *Aedes* mosquitoes and fruit flies, carboxylic acids are sensed by *Ir*-expressing neurons that we would denote as IRNs (Joseph & Carlson, 2015; Raji et al., 2019). It is therefore very likely that this acetic acid-sensitive sensillum houses *Ir*-expressing IRNs that are unaffected while the function of OR-expressing ORNs in adult chemosensation has been significantly compromised in *orco*^-/-^ *An. coluzzii*.

### Larval responses are largely but not exclusively Orco dependent

Most studies on the role of Orco in ORN olfactory transduction have focused on adult stages (Degennaro et al., 2013; Fandino et al., 2019; Larsson et al., 2004; Li et al., 2016), and few, if any, have examined its role in the pre-adult larval olfactory systems. In contrast to terrestrial adults, larval-stage mosquitoes are aquatic with a unique and robust chemical ecology as well as a distinctive set of chemosensory receptors (C. Liu et al., 2010; Xia et al., 2008). Recently, using wild-type *An. coluzzii* mosquitoes, we carried out a comprehensive *in vivo* electrophysiological characterization of the larval sensory cone, which is the primary olfactory organ for larval olfaction, detailing a robust and surprisingly complex set of olfactory responses (Sun et al., 2020). An examination of the electrophysiology of the *orco*^-/-^ mutant larval sensory cone revealed that, as expected, the overall background activity of sensory cone neurons was dramatically higher in the wild type (frequency of 45 ± 4 spikes/s) than in *orco*^-/-^ mutants (22 ± 3 spikes/s; Figure 5A). This supports previous studies indicating that the larval olfactory system of *An. coluzzii* uses both Orco-dependent ORNs as well as IRNs and perhaps other, Orco-independent transduction pathways (C. Liu et al., 2010). In contrast to the robust responses characteristic of wild-type larvae, *orco*^-/-^ mutant larvae were largely indifferent to nearly all (70 of 74) odorants in our test panel (Figure 5B). Nevertheless, *orco*^-/-^ mutant larvae still displayed responses to ammonia, 1-octanol, hexanoic acid, and 2,3-butanedione (Figure 5B, Supplementary Figure S3), as well as displaying inhibitory responses to a surprising number of odorants, such as 2,6-lutidine, 2-ethylphenol, pentylamine, 2-methyl-2-thiazole, when compared with activation in wild-type larval sensory cones (Figure 5B, Supplementary Figure S4).

### Reduced attraction to human-derived and human-mimic compounds in *An. coluzzii orco*^-/-^ mutant

Several odorants emanating from human skin comprise key attractants for host-seeking female mosquitoes, such as the highly anthropophilic *An. coluzzii*. In order to assess the relative contribution of Orco-expressing ORNs to those behaviors, we adapted a human-host proximity assay (Degennaro et al., 2013) that essentially counts the number of host-seeking female mosquitoes attracted to an unwashed human hand exposed within a chamber randomly attached to the side of a rearing chamber (Figure 6A). The studies revealed that while both wild-type and *orco*^-/-^ mutant host-seeking female *An. coluzzii* were indifferent to control (empty) chambers, wild-type individuals were robustly attracted to a human hand while *orco*^-/-^ females displayed significantly less attraction (Figure 6B). This suggests the absence of Orco expression in *An. coluzzii* significantly impairs the mosquito’s ability to robustly locate a human host. Host-seeking female Anopheline mosquitoes are also attracted to compounds thought to act as mimics of human-derived odors, such as Limburger cheese volatiles, which are presumptive foot-like odors (Knols & De Jong, 1996; Owino, 2011). To determine whether attraction to such human mimics relies on Orco-dependent pathways, we designed a simple two-choice behavioral bioassay to test the response of wild-type and *orco*^-/-^ mutant female *An. coluzzii* mosquitoes to Limburger cheese together with a heat source (for attraction and volatilization, Figure 6C). In these studies, while both wild-type and *orco*^-/-^ mutant females displayed similarly modest attraction to heat sources alone, the combination of heat and cheese attracted significantly more wild-type females than heat alone (Figure 6D). In contrast, host-seeking *orco*^-/-^ mutant females showed no difference between their attraction to a heat source with or without Limburger cheese augmentation (Figure 6D). These data are consistent with the hypothesis that Orco is required for the detection of and response to the human-mimic compound(s) in Limburger cheese volatiles.

**Figure 5.**
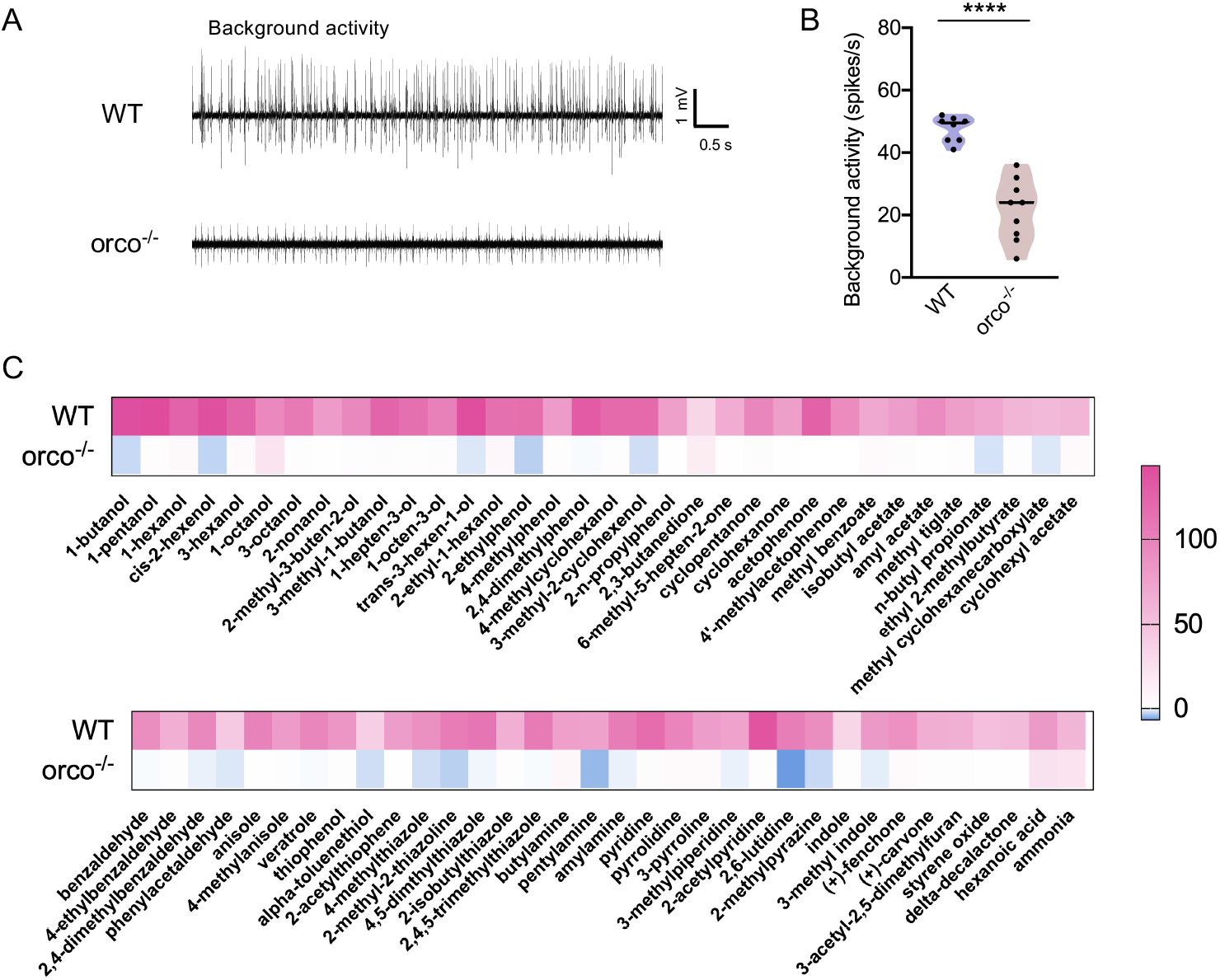
Orco-dependent response in the larval sensory cone. (A) Representative traces of single-unit electrophysiological comparison of the spontaneous (background) action potential spiking activity in the sensory core of wild-type and orco^-/-^ *An. coluzzii* larvae. (B) Collective background activity from wild-type and orco^-/-^ *An. coluzzii* larval sensory cones (n=6-8). (C) Heatmap presenting collective, normalized responses of the larval sensory cone of both wild-type and orco^-/-^ *An. coluzzii* to 74 odorant compounds. The wild-type *An. coluzzii* larval sensory cone data were originally reported in Sun et al., (2020).

**Figure 6.**
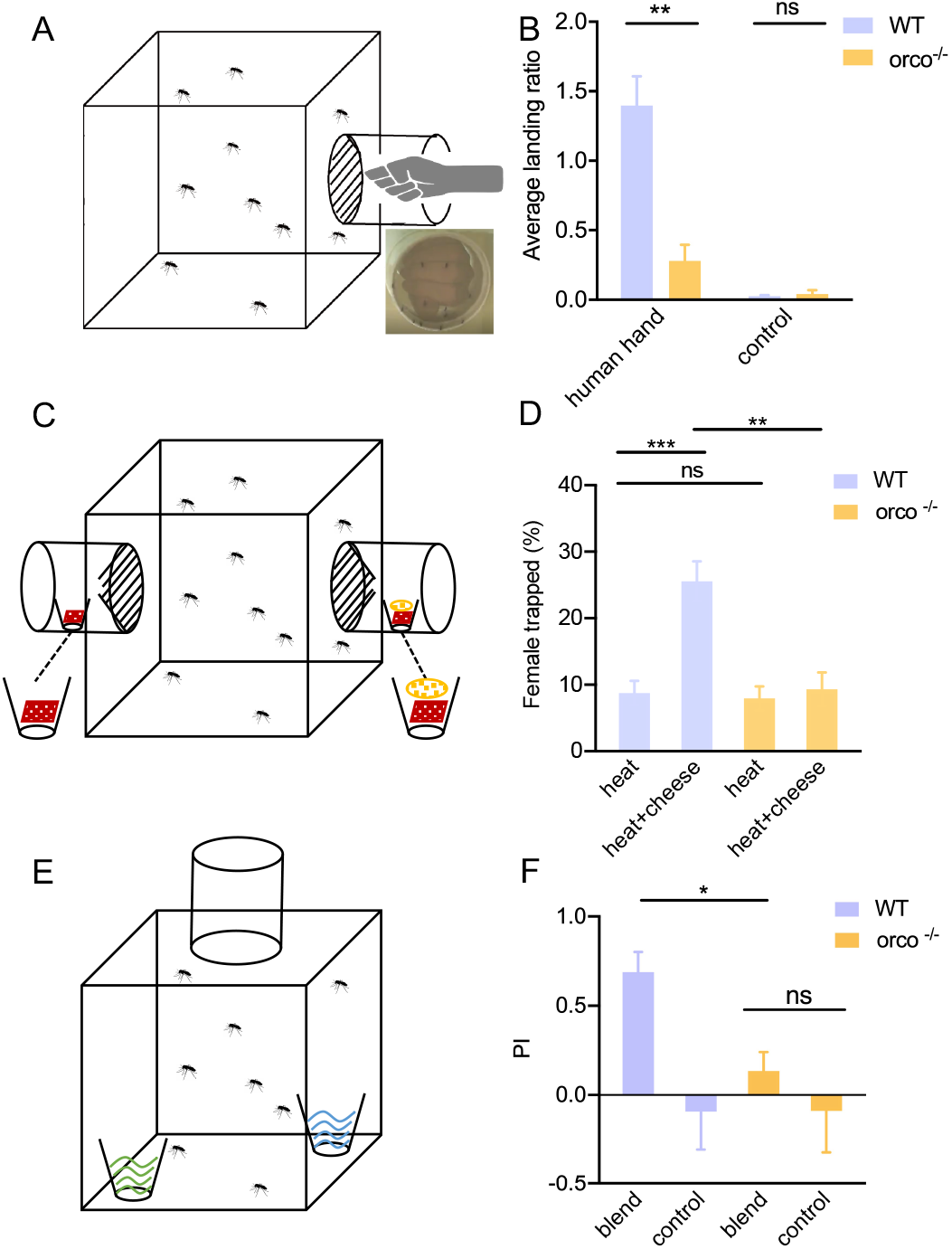
Behavioral responses of wild-type and orco^-/-^ mosquito to human- and non-human-derived odors: (C) Schematic graphic of the setup of human hand proximity bioassay; (D) Comparison of wild-type and orco^-/-^ mosquito hand attraction (n=7-11); (A) Schematic representation of the setup of the Limburger cheese human foot mimic bioassay; (B) Comparison of the wildtype and orco^-/-^ mosquito attraction to Limburger cheese to (n=6); (E) Schematic graphic of the setup of oviposition preference assay; (F) Comparison of attraction of gravid wild-type and orco^-/-^ female mosquitoes to the oviposition blend (n=5). Nonparametric Mann-Whitney test was applied in the statistical analysis, with p<0.05 (*), p<0.01 (**) and p<0.001 (***) indicating significant differences.

### Disruption of oviposition preference to an attractant blend

Oviposition preference by gravid female mosquitoes is also mediated to a large degree by their olfactory system (Montell & Zwiebel, 2016). To investigate the involvement of Orco-dependent processes in *An. coluzzii* oviposition behaviors, we next utilized a simple dual-choice oviposition assay (Figure 6E) along with a seven-compound blend (each component held at 10^−8^M) which has been reported as oviposition attractants for several mosquito species (Saveer et al., 2018). In these assays, gravid wild-type mosquitoes exhibited a robust preference relative to water+solvent-alone controls and oviposit in water cups supplemented with the attractant blend (Figure 6F).

However, this preference is not displayed by gravid *orco*^-/-^ mutant *An. coluzzii* females, which showed indistinguishably modest responses to the attractant blend as well as the water+solvent controls (Figure 6F). These data suggest that Orco is also directly involved in establishing the oviposition preference of gravid female *An. coluzzii*.

## Discussion

Mosquitoes utilize complex olfactory systems not only to locate and preferentially select hosts for blood meals but also to choose optimal sites for oviposition, and to avoid predators and other hazards. Inasmuch as the Orco co-receptor is universally required for the functionality of insect ORs, it is a central component of the mosquito’s olfactory signal transduction processes. We utilized CRISPR-CAS9-based gene targeting to generate an *orco*^-/-^ mutant in the malaria vector mosquito *An. coluzzii* to confirm that Orco is a critical component of the Anopheline olfactory system and is required for neuronal physiology as well as a range of downstream olfactory-driven behaviors. This aligns with previous findings in *Aedes* mosquitoes (Degennaro et al., 2013), in which *orco*-null mutants display dramatic deficiencies in ORN processes as well as a range of behavioral deficits, including alterations in host seeking.

In this study, we have characterized *An. coluzzii* Orco functionality *in vivo* in both adult and larval olfactory transduction pathways, where it clearly plays an essential role in a number of cell types and contexts. These include the larval sensory cone ORNs as well as the adult maxillary palp and antennae where cpB/C ORNs as well as the neurons associated with the vast majority of trichoid sensilla show profound deficits in background and odorant-induced signaling (Figure 4). These data also confirm the Orco-independent functionality of Anopheline maxillary palp cpA neurons, which remain robustly responsive to CO_2_ as well as a distinct subset of larval sensory cone neurons and adult antennal trichoid sensilla tuned to acids and amines such as ammonia. These differences affirm current paradigms in which diverse chemosensory neurons spread throughout adult and larval olfactory appendages express an assortment of OR, GR and IR cell surface receptors that together comprise a mosaic array that provides an expansive capacity for combinatorial odor coding required for complex responses to environmental cues and other stimuli.

*An. coluzzii orco*^-/-^ mutants also display a range of behavioral phenotypes. As observed in *Ae. aegypti* mosquitoes (Degennaro et al., 2013), direct attraction of blood-feeding females to unwashed human hands as a paradigm for host seeking was severely impacted in our *orco*^-/-^ mutants (Figure 6A, B). We also used Limburger cheese, which is a well-established human-odor proxy for anthropophilic Anophelines (Knols & De Jong, 1996; Owino, 2011); when coupled to a heat source, these mimics strongly attracted wild-type mosquitoes while *orco*^-/-^mutants were indifferent (Figure 6C, D). GC-MS studies have identified multiple Limburger cheese volatiles, including ketones (e.g. 2-pentatone, 2-heptanone, 2-octanone, and 2-nonanone) and phenols (e.g. phenol, phenylethanol and cresol) (Bertuzzi et al., 2018), that are also detected in human skin emanations (Bernier et al., 2000). Indeed, many of these odorants have been shown to directly and robustly activate multiple Anopheles Orco/Or complexes (Carey *et al*., 2010; Wang *al*., 2010), making it altogether likely that these compounds act as human blood-meal host cues that normally activate a set of ORNs that are effectively silenced in *An. coluzzii orco*^-/-^ mutants. We also found that gravid *An. coluzzii orco*^-/-^ mutants are behaviorally insensitive to blends of oviposition attractants that strongly attract wild-type gravid mosquitoes (Figure 6E, F). Once again, several components of these blends have been shown to directly activate multiple ORs when functionally expressed with an Orco ortholog (Carey *et al*., 2010; Wang *et al*., 2010). For example, 2-propylphenol, which attracts gravid *An*. colluzzi seeking oviposition sites (Rinker et al., 2013), evokes a strong response in Drosophila transgenically expressing *An. coluzzii* Or9 (Carey et al., 2010).

We recently completed a comprehensive characterization of the larval sensory cone in wild-type *An. coluzzii* (Sun et al., 2020). Thus, in addition to analyzing the phenotypes of adult *orco*^-/-^ mosquitoes, we were also able to explore how this mutation affected the physiology of the larval sensory cone, which is the primary larval olfactory appendage. While background activity and odor-evoked responses were considerably lower, the *orco*^-/-^ larval sensory cone nevertheless displayed some interesting features of residual activity. This included significantly less background activity across the sensory cone as well as uncovering modest and, in some cases, pronounced inhibition in response to several odorants that normally activate the larval sensory cone. It is likely that the lower overall background spiking is due to the silencing of Orco-expressing ORNs while the collective inhibition in response to odorant stimulation remains enigmatic. In any case, these inhibitory effects occur while a discrete activation paradigm of the larval sensory cone in response to ammonia and hexanoic acid remains largely unperturbed. The ability of these neurons to continue firing at near-wild-type levels suggests the presence of dendrites derived from other types of chemosensory neurons in addition to those associated with *Orco*-expressing ORNs. Indeed, previous studies have identified multiple IRs in larval antennae (Liu et al., 2010), making it very likely that this residual sensory cone activity in *orco*^-/-^ larvae originates from IR-expressing IRNs. Coeloconic sensilla of adult Drosophila antennae, which exclusively house IRNs, robustly respond to 1-octanol, hexanoic acid and 2.3-butanedione (Benton et al., 2009), the same three compounds that maintain mild stimulation of the *An. coluzzii orco*^-/-^ larval sensory cone (Figure 5C). It is therefore reasonable to assign this activity to *Ir*-expressing IRNs, which send dendrites to populate the larval sensory cone and act independently of *orco* expression and functionality in larval ORNs.

These studies confirm that, as is the case for a growing list of insect species, Anopheline mosquitoes require a functional copy of the Orco co-receptor in order to carry out a wide range of olfactory processes. In contrast to the large number of highly diverse and differentially expressed tuning ORs in *An. coluzzii* and in other insects, the importance of *Orco* is underscored by its broad expression across insect ORNs as well as its extreme sequence conservation and functional orthology. While the detection and responses to diverse sensory modalities, such as thermal, visual and mechanosensory cues, as well as a complementary suite of Orco-independent chemosensory pathways provide phenotypic plasticity that facilitates the survival of *orco*^-/-^ mosquitoes in the laboratory, it is likely that the loss of *orco* functionality would create a lethal disadvantage in nature. As such and in light of its unique conservation across insect taxa, targeting *orco* for the design of novel paradigms for the control of insects that act as disease vectors as well as a range of agricultural and other pests remains an alluring prospect.

---

**Table S1:**
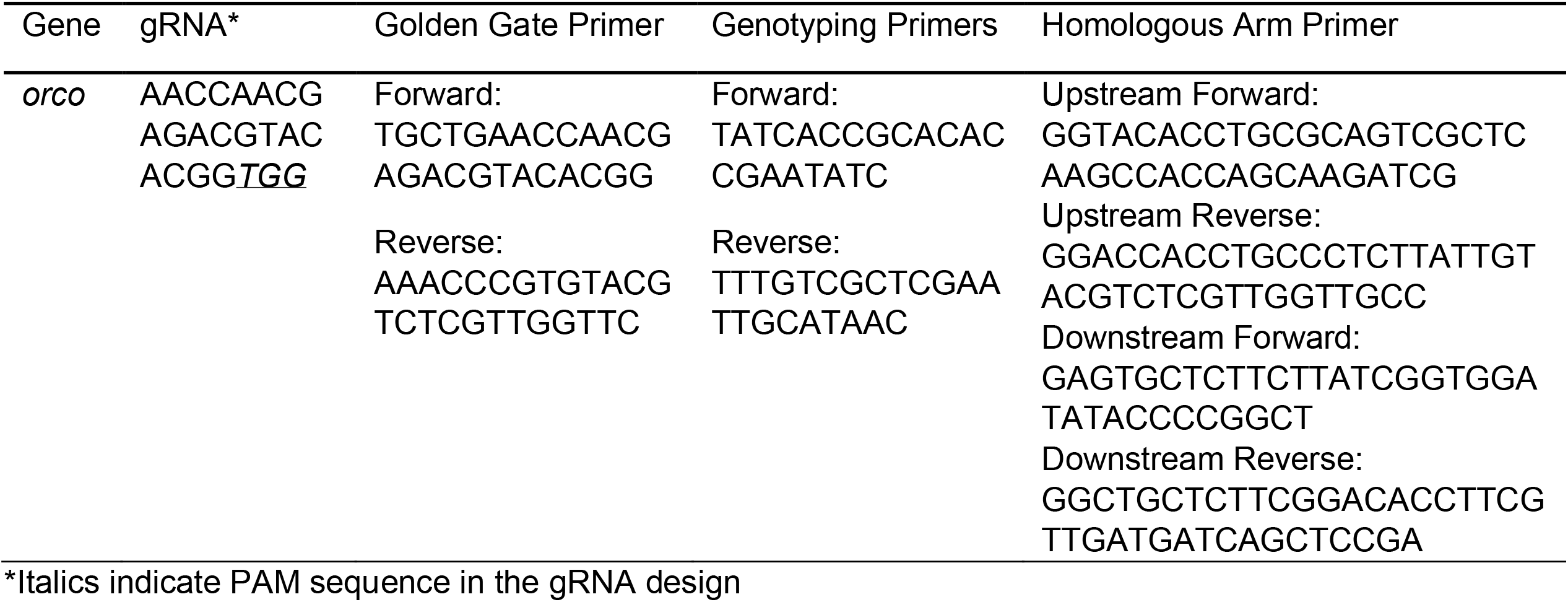
Oligonucleotides used for generating the *orco* gene-mutated mosquitoes.

**Table S2.** Normalized EAG responses of WT and orco^-/-^ *An. coluzzii*.

|  | Orco <sup>-/-</sup> |  | WT |  |
| --- | --- | --- | --- | --- |
|  | Mean | SEM | Mean | SEM |
| isoamyl acetate | 1.19 *** | ± 0.14 | 3.14 | ± 0.15 |
| isobutyl acetate | 0.68 *** | ± 0.07 | 2.67 | ± 0.18 |
| ethyl propionate | 0.83 *** | ± 0.11 | 2.10 | ± 0.19 |
| geranyl acetate | 1.75 | ± 0.20 | 2.78 | ± 0.36 |
| ethyl butyrate | 0.85 *** | ± 0.10 | 2.29 | ± 0.15 |
| ethyl acrylate | 0.76 * | ± 0.21 | 2.14 | ± 0.31 |
| methyl salicylate | 0.46 *** | ± 0.09 | 2.17 | ± 0.44 |
| benzyl acetate | 1.38 * | ± 0.09 | 2.25 | ± 0.35 |
| 6-methyl-5-hepten-2-one | 0.71 ** | ± 0.12 | 2.03 | ± 0.33 |
| 2-nonanone | 1.49 ** | ± 0.23 | 6.06 | ± 0.78 |
| 3-octanone | 1.55 ** | ± 0.32 | 4.51 | ± 0.39 |
| 2,3-hexanedione | 1.66 ** | ± 0.27 | 5.52 | ± 0.73 |
| 3-pentanone | 0.95 ** | ± 0.22 | 4.71 | ± 0.98 |
| 2,3-pentanedione | 1.70 ** | ± 0.35 | 4.42 | ± 0.71 |
| 2'-hydroxyacetophenone | 0.43 *** | ± 0.04 | 2.76 | ± 0.46 |
| 2-isopropylphenol | 1.38 | ± 0.42 | 4.14 | ± 1.49 |
| 1-hexen-3-ol | 0.92 ** | ± 0.12 | 4.34 | ± 0.89 |
| 1-pentanol | 0.77 ** | ± 0.16 | 6.18 | ± 1.15 |
| 1-butanol | 0.41 ** | ± 0.22 | 5.81 | ± 0.99 |
| (s)-1-octen-3-ol | 0.64 ** | ± 0.12 | 4.22 | ± 0.39 |
| 3-hexanol | 0.57 ** | ± 0.11 | 4.60 | ± 0.51 |
| hexanal | 1.25 ** | ± 0.24 | 3.85 | ± 0.53 |
| acetal | 1.61 ** | ± 0.13 | 2.66 | ± 0.21 |
| octanal | 1.51 *** | ± 0.12 | 4.34 | ± 0.35 |
| nonanal | 1.52 *** | ± 0.12 | 3.87 | ± 0.49 |
| 4-isopropylbenzaldehyde | 1.32 * | ± 0.16 | 2.20 | ± 0.33 |
| (+)-fenchone | 1.28 ** | ± 0.23 | 4.57 | ± 0.65 |
| linalool oxide | 0.48 ** | ± 0.11 | 5.85 | ± 0.67 |
| (DL)-camphor | 1.07 *** | ± 0.10 | 2.33 | ± 0.15 |
| 2-acetylthiophene | 1.26 * | ± 0.29 | 2.77 | ± 0.41 |
| 2-acetylthiazole | 1.12 * | ± 0.08 | 2.29 | ± 0.34 |
| 2,4-dimethylthiazole | 0.49 ** | ± 0.09 | 6.55 | ± 1.06 |
| 2,4,5-trimethylthiazole | 1.18 *** | ± 0.14 | 3.10 | ± 0.16 |
| 2-acetylpyridine | 0.63 ** | ± 0.12 | 4.76 | ± 0.50 |
| butylamine | 2.12 | ± 0.55 | 2.56 | ± 0.56 |
| 3-methylpiperidine | 8.16 | ± 1.13 | 7.68 | ± 1.33 |
| pyrrolidine | 5.27 ** | ± 0.52 | 3.18 | ± 0.58 |
| 2-oxobutyric acid | 3.29 | ± 0.54 | 3.57 | ± 0.37 |
| L-(+)-lactic acid | 3.56 | ± 0.82 | 2.23 | ± 0.27 |

Nonparametric Mann-Whitney test was applied for statistical analyses; significant differences indicated by p<0.05 (\*), p<0.01 (\*\*) and p<0.001 (\*\*\*)

## Figure legends

**Supplemental Figure S1.**
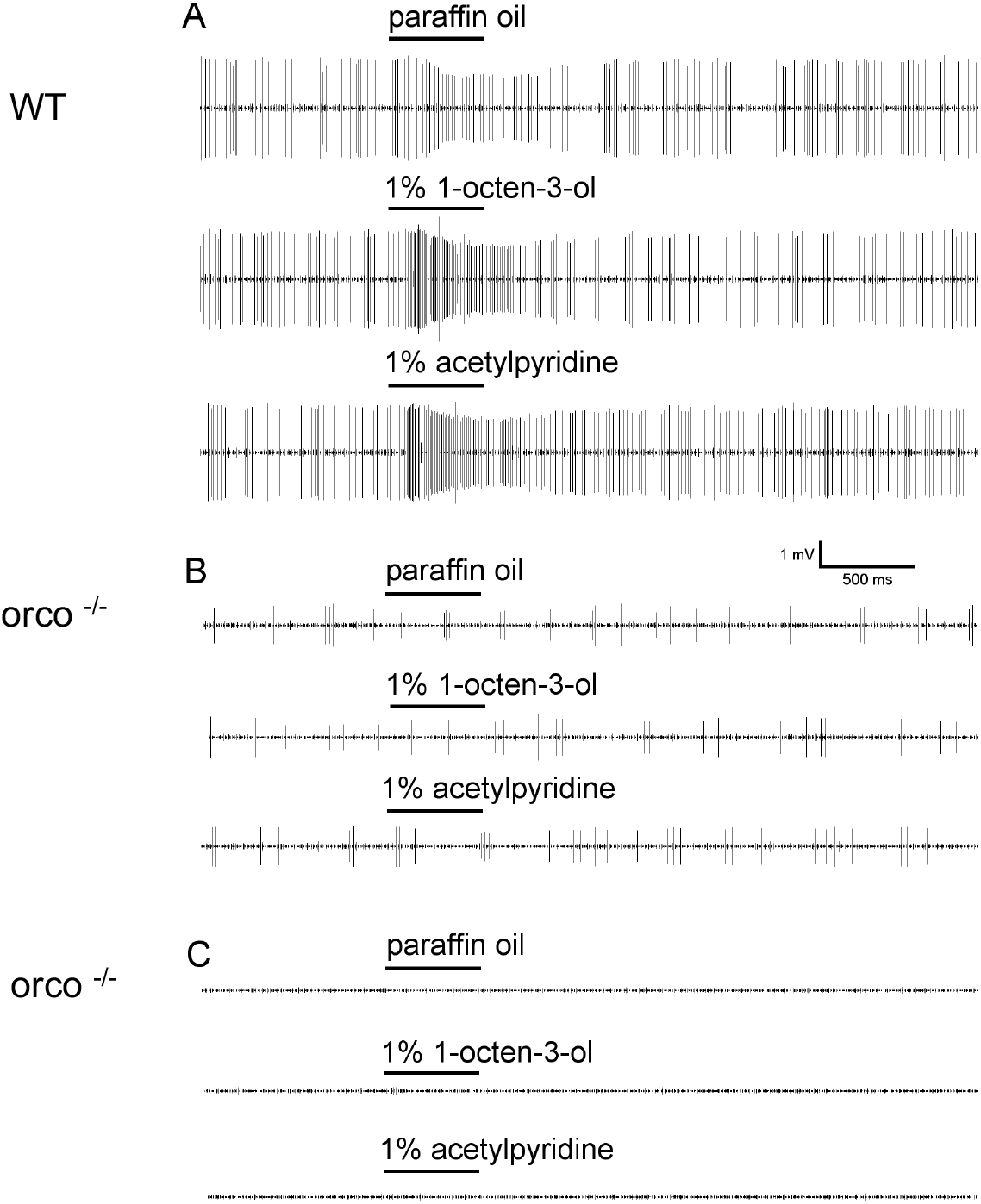
Representative response traces of trichoid sensilla to odors: (A) Trichoid sensilla of wild-type mosquitoes in response to paraffin oil (solvent), 1-octen-3-ol, 2-acetylpyridine at 100-fold dilution; (B) Trichoid sensilla of *orco* mutant mosquitoes in response to paraffin oil (solvent), 1-octen-3-ol, 2-acetylpyridine at 100-fold dilution with residue spike; (C) Trichoid sensilla of *orco* mutant mosquitoes in response to paraffin oil (solvent), 1-octen-3-ol, 2-acetylpyridine at 100-fold dilution without residue spike.

**Supplemental Figure S2.**
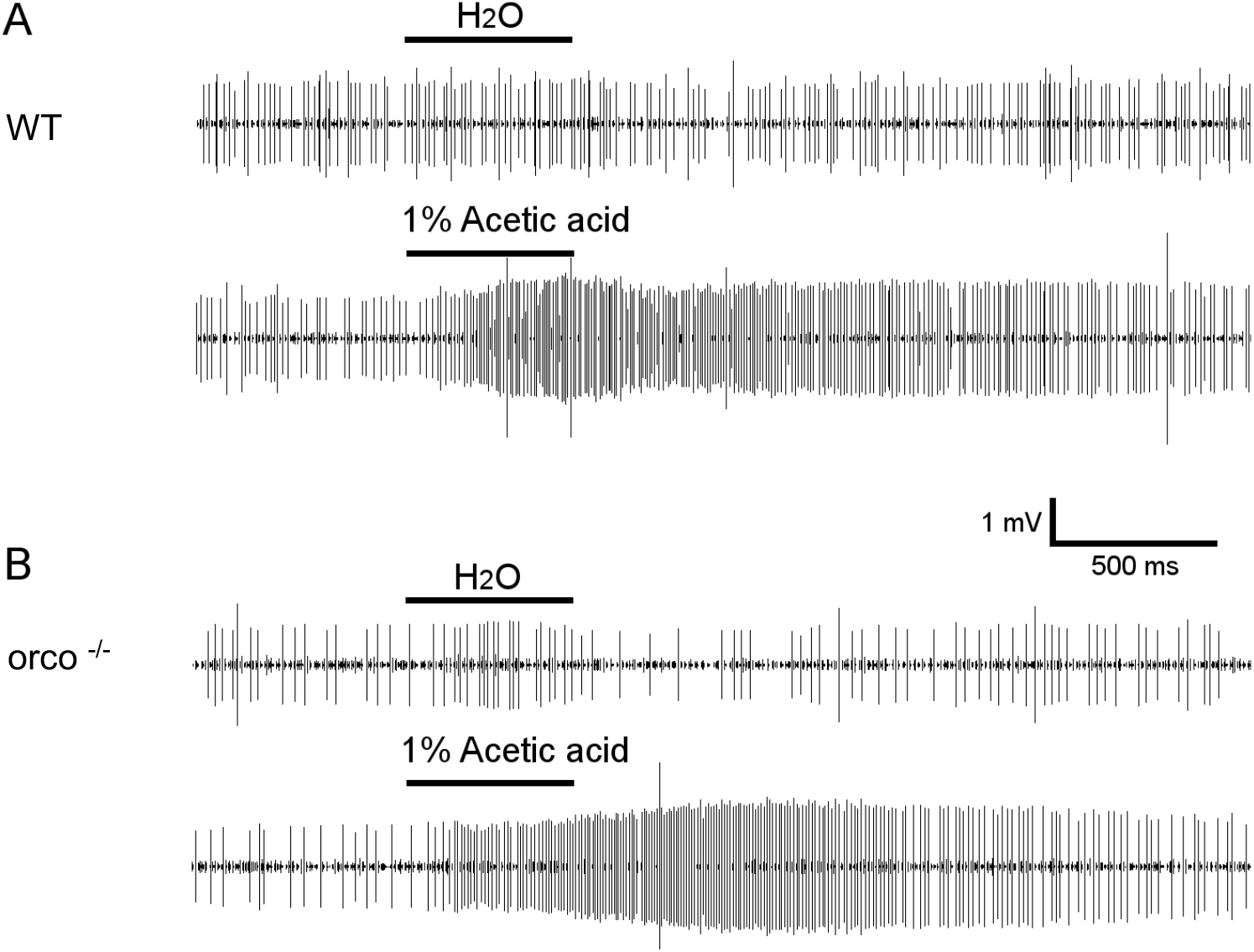
(A) Representative response traces of acid-sensitive trichoid sensilla in the wild-type mosquito challenged with H_2_O (solvent) and 1% acetic acid; (B) Representative response traces of acid-sensitive trichoid sensilla in orco^-/-^ mosquitoes challenged with H_2_O (solvent) and 1% acetic acid.

**Supplemental Figure S3.**
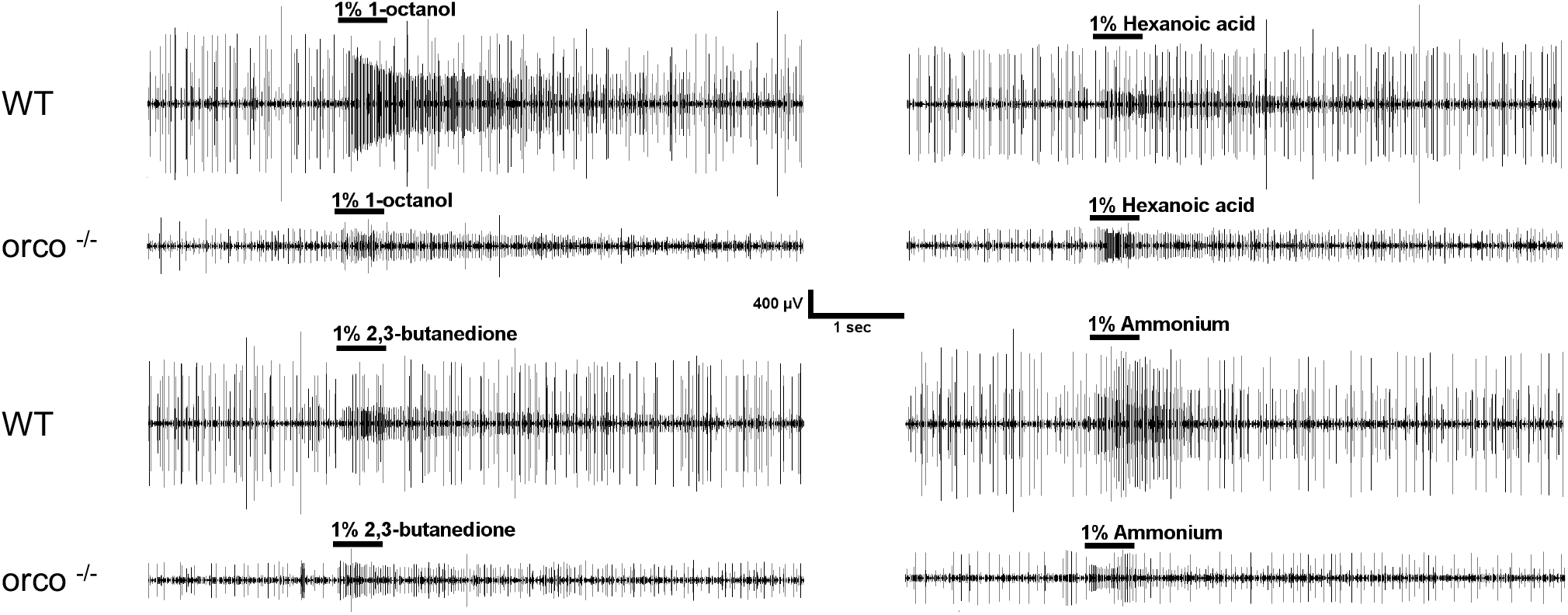
Representative response traces of larva sensory cone of both wild-type and orco^-/-^ *An. coluzzii* to multiple odorants eliciting excitatory neuronal firing.

**Supplemental Figure S4.**
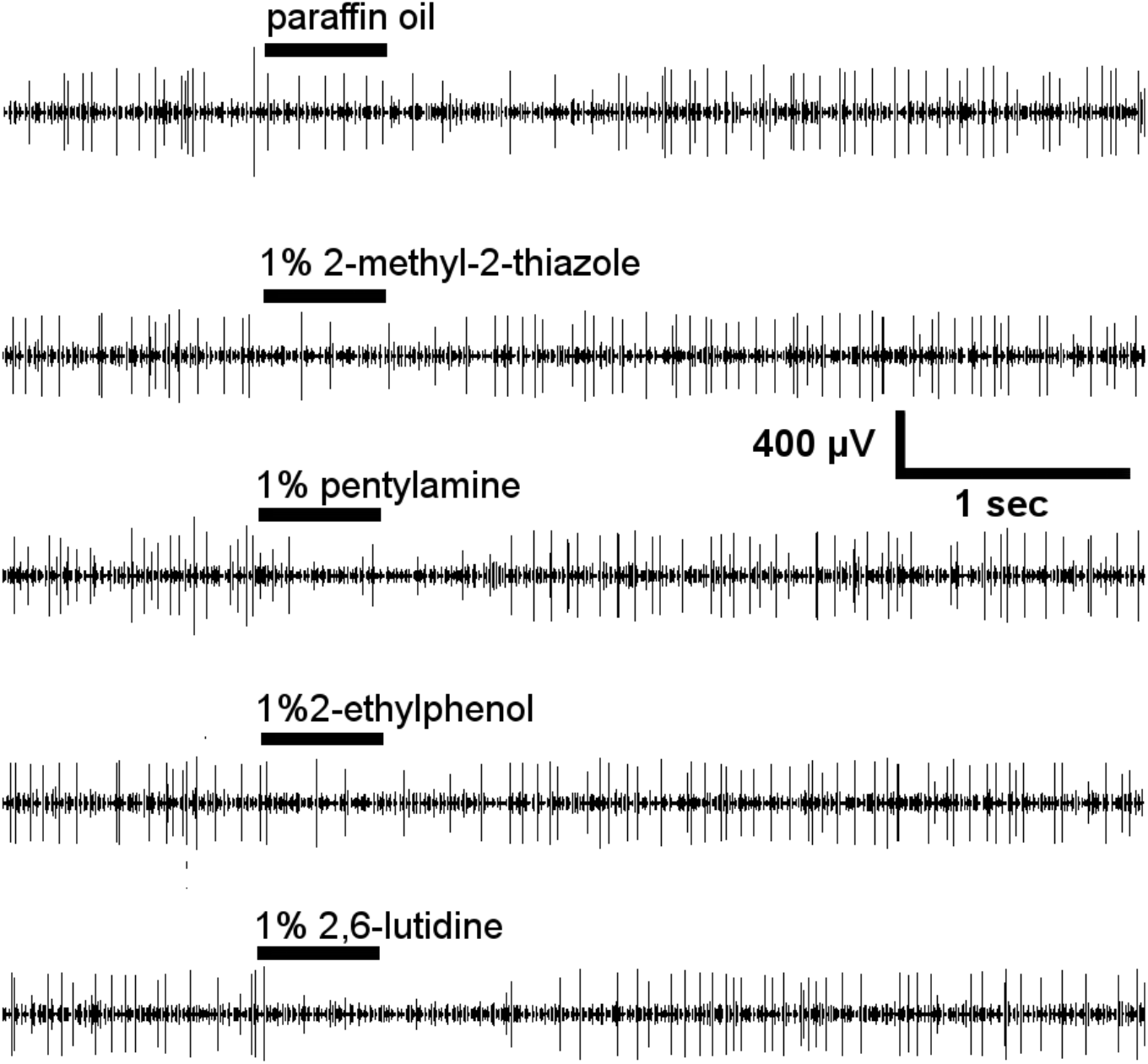
Representative response traces of larva sensory cone of orco^-/-^ *An. coluzzii* to multiple odorants eliciting inhibitory neuronal firing.

## Acknowledgments

We wish to thank Dr. Andrea Crisanti of Imperial College London for the CRISPR gene targeting vector and Dr. H. Willi Honegger, Dr. Ann Carr, Stephen Ferguson and other members of the Zwiebel lab for their comments on this manuscript, critical suggestions and help with data analyses during the course of this work. We also thank Zhen Li and Samuel Ochieng for mosquito rearing and technical help. This work was conducted with the support of Vanderbilt University and a grant from the National Institutes of Health (NIAID, AI127693) to LJZ.

